# Unwilling or Unable? Using 3D tracking to evaluate dogs’ reactions to differing human intentions

**DOI:** 10.1101/2022.07.09.499322

**Authors:** Christoph J. Völter, Lucrezia Lonardo, Maud G.G.M. Steinmann, Carolina Frizzo Ramos, Karoline Gerwisch, Monique-Theres Schranz, Iris Dobernig, Ludwig Huber

## Abstract

The extent to which dogs (*Canis familiaris*) as a domesticated species understand human intentions is still a matter of debate. The unwilling-unable paradigm has been developed to examine whether nonhuman animals are sensitive to intentions underlying human actions. In this paradigm, subjects tended to show more patience toward a human that appears willing but unable to transfer food to them compared to an unwilling (teasing) human. In the present study, we conducted the unwilling-unable paradigm with dogs using a detailed behavioural analysis based on machine-learning driven 3D tracking. Throughout two preregistered experiments, we found evidence, in line with our prediction, that dogs reacted more impatiently to actions signalling unwillingness to transfer food rather than inability. These differences were consistent through two different samples of pet dogs (total N=96) and they were evident also in the machine-learning generated 3D tracking data. Our results, therefore, provide robust evidence that dogs distinguish between similar actions (leading to the same outcome) associated with different intentions. However, their reactions did not lead to any measurable preference for one experimenter over the other in a subsequent transfer phase. We discuss different cognitive mechanisms that might underlie dogs’ performance in this paradigm.

## Introduction

Interpreting others‘ actions in terms of goals and intentions is fundamental for us humans to make sense of our social environment [1]. Intention has been defined as an action plan to achieve a certain outcome, the goal [2,3]. Understanding intentions allows us to react appropriately to others’ actions, predict impending actions and interact and communicate with others effectively.

In the case of successful intentional actions — for instance an actor has the goal of opening a box and does so — the methodological challenge for studying intention reading is that the actor’s goal and the environmental outcome match [4]. Therefore, it is impossible to disentangle whether the subject reacts to the result of the demonstrator’s action or her/his intention. To study intention reading (i.e. the understanding of intentions that leading to specific actions), researchers have created situations in which similar actions imply different intentions, contextual information leads to different interpretations with respect to the intentions of an action, or incomplete (failed) actions are shown but the intention can nevertheless be inferred [4–6]. In these cases, one can then compare the subjects’ responses to failed attempts, accidents, impossible tasks and deliberately unsuccessful attempts, all of which lead to the same external outcome but are underscored by different intentions.

The understanding of others’ intentions seems to develop in humans towards the end of the first year of life. For example, Behne et al. [5] compared infants’ reactions to an experimenter that was unwilling to transfer a toy to the infant (i.e., the experimenter teased the infant or played with the toy on her own) with an experimenter that was willing but unable to transfer the toy (i.e., she clumsily dropped the toy while trying to pass it on to the infant). Infants from the age of 9 months reacted more impatiently toward the unwilling than unable experimenter (evidenced by higher rates of reaching toward the toy and looking away from the experimenter). Today there is ample evidence that preverbal human children are able to infer the intentions that underlie actions and communicative signals based on subtle behavioural differences between intentional and accidental actions [5,7], failed actions [6] and situational constraints [8,9].

Understanding intentions might also be beneficial for nonhuman animals because it might allow them to interact effectively with social partners and competitors. Indeed, comparative evidence is consistent with the notion that we share the ability to understand intentions at least with some other species: using the same unwilling-unable paradigm, Call and colleagues (2004) investigated how zoo-housed chimpanzees (*Pan troglodytes*; N=12) responded to a human that acted as if either unwilling or unable to transfer food to them. In this study, the human experimenter would hand pieces of fruit to the subject through a small opening in a Plexiglas panel. Within this feeding routine, some test trials were interspersed in which the experimenter withheld the food. In some of these trials (the unwilling condition), the experimenter did so intentionally: for example, the experimenter would perform a teasing movement bringing food close to the subject but then retracting it in a teasing manner before the chimpanzee could access it. In other test trials (the unable condition), the experimenter intended to hand food to the subject, but for different reasons (e.g., clumsily dropping the food; the feeding hole being too small) the experimenter failed to do so. The chimpanzees produced begging and assertive behaviours more frequently when the experimenter was teasing compared to when a situational constraints prevented the experimenter from passing the food on to the subject (no significant differences were found between the teasing and clumsy condition with respect to the begging/assertive behaviours). The chimpanzees were also significantly faster in leaving the testing station in the teasing condition than the clumsy condition (no significant differences were found between the teasing and blocked condition). Similar findings (differing reactions to a teasing or clumsy experimenter) were reported for capuchin monkeys (*Sapajus paella*; N=6 [10]) and horses (*Equus caballus*; N=21 [11]).

Tonkean macaques (*Macaca tonkeana*; N=15 [12]) and African grey parrots (*Psittacus erithacus*; N=3; [13]) showed differing reactions between the teasing and blocked conditions; the clumsy condition was not administered in the latter two studies.

Dogs constitute a particularly interesting species with respect to reading human intentions as they have lived alongside humans for thousands of years. Dogs are known to closely monitor human actions, communicative signals, and facial expressions [14,15] and there is evidence that dogs’ cognitive abilities have adapted to living in human society [16,17]. For example, dogs understand human pointing gestures in a way other species (such as chimpanzees) do not [18]. They are also sensitive to ostensive cues in this context [19–23]. However, there is controversy around the cognitive abilities involved. Do dogs understand intentions underlying human actions or have they learnt a set of behavioural rules that link certain behaviours and outcomes? While the latter option was the predominant view for a long time [14,24], there is accumulating evidence of the flexibility and complexity of dogs’ reactions to human actions and attentional states [25–30].

For instance, there is evidence that dogs attribute goals to human agents. The benchmark test for goal-based action understanding is a paradigm established by Woodward [31] with human infants.

Woodward habituated infants to a hand moving across a stage and grasping one of two objects. After the habituation, the positions of the two objects were reversed and one of two new events was shown: either the hand took the same path to grasp a different object or it took a different path to grasp the previously chosen object. Infants looked longer at the new-goal/old-path event. Woodward’s findings suggested that infants encoded the actor’s action in relation to its goal. Presented with the same paradigm, dogs showed a similar looking time pattern [28] when watching a human agent approaching and touching one of two objects but not when an inanimate object performed the same actions (a black box approaching one of the goal objects).

Another piece of evidence for goal attribution in dogs comes from studies with do-as-I-do trained dogs. In one study, a dog completed the human demonstrator’s mimed action of hurdle jump (actually without a hurdle) by jumping across a hurdle that was located at some distance from the human’s jumping location [32]. In another study, dogs engaged in emulation (copying of the outcome) of an object-related action more often if the demonstration included an obvious action goal, whereas they showed a higher degree of copying fidelity when the action goal was unclear [33]. Finally, there is also evidence that dogs distinguish between intentional and accidental actions. Dogs followed a pointing gesture more frequently when the experimenter made eye-contact (an ostensive signal) than when she looked at her watch [19] highlighting the importance of ostensive signal for dogs [22,23,34].

A recent study found evidence that also dogs (N=51) distinguished between human actions signalling unwillingness or inability [35]. In contrast to the previous unwilling-unable studies, the dogs could walk around the barrier to approach the experimenter directly (the dogs were familiarised with this response prior to the test). The main finding was that the dogs walked around the barrier to approach the experimenter significantly sooner in the unable blocked and clumsy conditions than in the unwilling (teasing) condition. However, the difference in the setup makes the findings difficult to compare to the previous studies: The food dropped on the floor in the clumsy condition might have motivated the dogs to approach the experimenter given that, in their daily lives, they presumably often obtain food dropped on the floor. Moreover, the experimenter also produced different utterances in the test trials in contrast to previous studies (teasing: “ha-ha!”, clumsy: “oops!”, blocked: “oh!”) increasing the risk that the dogs differing reactions were driven by associations learnt prior to the test or by different tones of voice across conditions. In the current study, we therefore administered the unwilling-unable paradigm with dogs in a more consistent manner compared to the existing literature.

In the current pre-registered study, our goal was three-fold: first, we examined whether dogs indeed differentiate between actions signalling unwillingness vs. inability even if they cannot approach the experimenter or the dropped food (clumsy condition) directly (cf. [35]). Second, we investigated whether this distinction could also be identified without a human observer manually scoring the videos using machine-learning driven 3D tracking [36]. This served to mitigate the risk of inadvertent biases of the human raters. Third, in a follow-up experiment we explored whether the dogs’ differing reaction would transfer to subsequent preference and point-following tasks. Based on the previous findings [35], we hypothesized that dogs would be inclined to wait more patiently for a piece of food when the experimenter was unable rather than unwilling to pass it to them. Hence, we predicted that the dogs would spend more time away from the barrier separating them from the experimenter, sit or lay down for a longer period, and look away from the experimenter more frequently in the unwilling-teasing condition especially compared to the two unable-clumsy condition. With respect to the unable-blocked condition, our prediction was less clear because we predicted that the dogs might understand the inaccessibility of the reward due to the added blockage.

## Experiment 1

### Methods

#### Subjects

Forty-eight pet dogs of various breeds and mixed-breeds participated in this study (mean age: 77.5 months; range: 13 and 158 months; 31 females and 17 males). Seven additional dogs were tested but excluded because they were too young (<10 months; N=3) or because they did not take the food directly from the experimenter (N=4). The sample size was preregistered and based on a previous unwilling-unable study with dogs [35]. The study was discussed and approved by the institutional ethics and animal welfare committee in accordance with GSP guidelines and national legislation (approval number: ETK-028/02/2021).

#### Materials

The study took place in one of the test rooms (6m x 7m) of the Clever Dog Lab, Vienna. During the test, the experimenter (E) and the subject were separated by a barrier consisting of a central, transparent polycarbonate panel inside a stable wooden frame (w x h: 120 × 110.5cm) and two lateral wire-mesh fences (w x h: 179.5 × 107.3cm) perpendicular to the panel (Figure 1A). The U-shaped barrier was located such that lateral fences were in contact with a wall of the room such that the fenced area was closed off and the subjects could not enter it.

**Figure 1.**
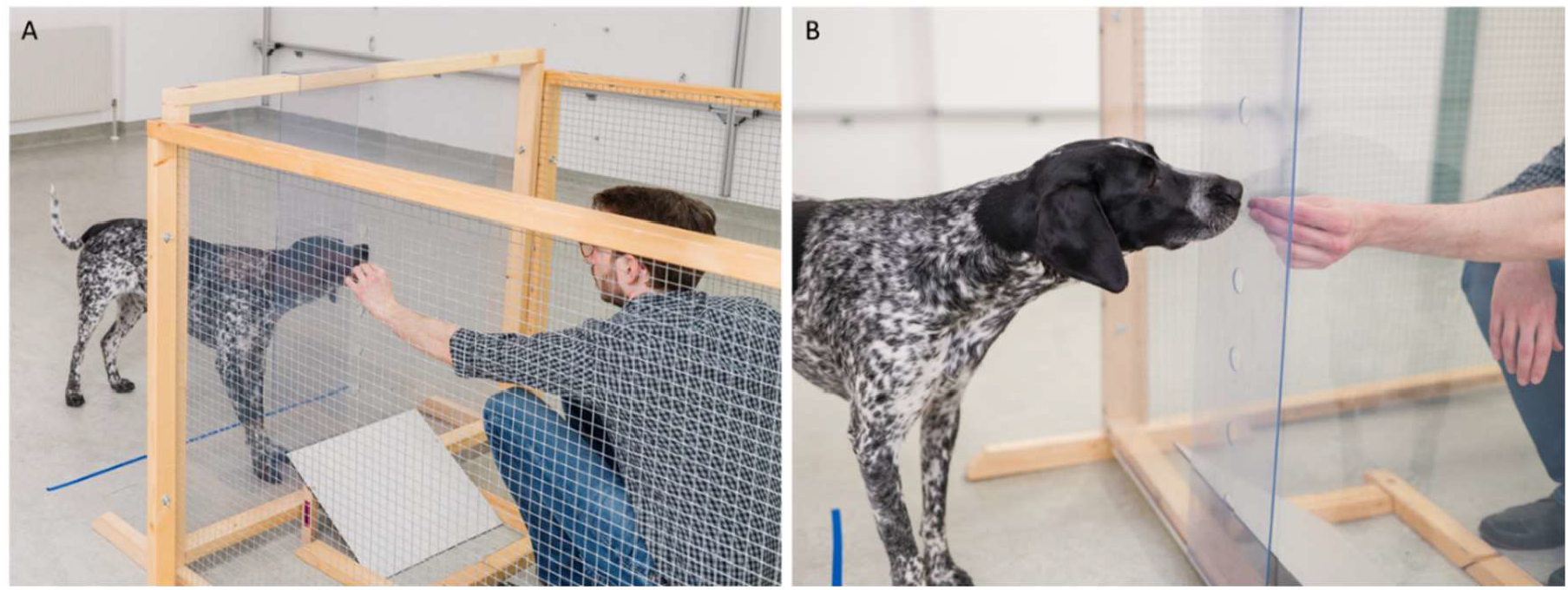
Experimental setup. A. The experimenter gives food reward to the dog through a hole in the polycarbonate panel during a motivation trial. B. The holes in the panel are covered by a polycarbonate panel during the blocked condition. Informed consent was obtained for publication of identifying images (photo credits: Thomas Suchanek/Vetmeduni).

In the centre of the polycarbonate panel were six small holes (diameter: 3cm) positioned at different heights. The lowest hole was at 20.5cm measured from the floor, the highest at 70.5cm, with 10cm gaps between the holes. During the experiment, E sat on a small stool inside the fenced area facing the polycarbonate panel with the holes inside. E wore an FFP2 mask throughout the experiment. Underneath the polycarbonate panel, we placed a wooden ramp (w x d x h: 35 × 45 × 20cm) in such a way that anything that fell on the surface, rolled away from the polycarbonate panel. Wooden dowels on the floor in between the stool and the ramp ensured that the food items that rolled down the ramp would not roll away. Attached to the outside of the polycarbonate panel was another piece of polycarbonate (w x h: 24 × 110cm) that the experimenter could slide in front of the holes to block them (Figure 1B). We added a line of coloured tape to the floor 50 cm away from the panel on the dog’s side, which served as a marking for the behavioural scoring. The dogs’ caregiver sat on a chair located at the wall opposite to the fences area (ca. 3.5m away from the fenced area). The dogs were allowed to move freely in the room during the study.

#### Design

We conducted a within-subject design with three conditions: the teasing, clumsy, and blocked condition (Video S1). In all three conditions, the (same) E brought a food reward (e.g. a slice of sausage) close to a hole in the polycarbonate panel that separated dog and E but never actually transferred the food to the subject. In the teasing condition, the E quickly pulled the food back when it reached the hole in the panel in a teasing-like manner. In the clumsy condition, she dropped the food once it reached the hole in the panel. The food fell on the ramp underneath and rolled back to E. In the blocked condition, an additional Polycarbonate segment blocked the holes.

Each subject completed 12 test trials, 4 trials per condition (with motivation trials before and in between the test trials). The conditions were presented in blocks. The order of the conditions within the test session were counterbalanced across subjects. In the motivation trials, E transferred the food successfully and immediately to the dog. At the beginning of each block, there were four motivation trials and then two motivation trials interspersed after each test trial. Hence, the dog got food more often than not, to maintain their motivation. After each block, there was a short break.

#### Procedure

At the beginning of the session, the caregiver entered the room with the dog, sat down on the chair opposite to the experimental setup and released the dog. The caregiver was instructed to interact with the dog during the session only on a signal by E. Then E entered the room and the fenced area and sat down on the stool. E called the dog with a piece of sausage in her hand. Once dog approached the panel and took the offered food, E started the next motivation trial by retrieving another piece of sausage from a container behind her back.

Before the start of each test trial, E raised her hand to signal to the caregiver to call the dog back. While the dog walked back to the caregiver, in the blocked condition, E blocked the holes in the panel by sliding the blocker in front of the holes (unseen by the dog). At the start of a test trial, E offered another piece of sausage to the dog and started the demonstration once the dog had crossed a marking on the floor 50cm away from the panel. The demonstrations took 30 seconds. In all test conditions, E moved the food forward to one of the holes in the panel. We used the same hole throughout the session but adjusted the height to each dog’s shoulder height. In the teasing condition, E moved the sausage back abruptly and then moved it forward again (in a teasing manner). In the clumsy, condition E dropped the food right at the hole in the panel. The food fell down on the ramp underneath and rolled away from the panel toward E. E then picked it up and moved it forward again. In the blocked condition, E moved the food forward until it touched the polycarbonate piece blocking the holes and turned the hand left and right as if pushing the food through a narrow opening. E then moved food back in the same way as in teasing condition. In the test trials, E repeated the unsuccessful food transfer attempts until the predetermined trial duration of 30 s was over (the 30-s period started when the dog crossed the 50-cm-line in front of the panel). E continued executing the actions irrespective of whether the dog stayed at the panel. E maintained a neutral facial expression and did not communicate verbally with the subjects during the test trial.

#### Scoring

The test was videotaped with two separate camera systems (see supplementary material). For the behavioural scoring, we used the video analysis software Loopy (http://loopb.io, loopbio Gmbh, Vienna, Austria). We scored the durations the dogs spent sitting or lying down (resting), the durations the dogs spent with all feet further away than 50cm from the polycarbonate panel (time away), the durations the dogs approached the fenced area from the side (lateral approach), and whether the dogs looked away from the experimenter/barrier. We scored lateral approach when the dogs were in close proximity of the fence segments and their head was oriented to the fence. We scored looking away when the dog’s snout was directed away from the barrier such that it was approximately parallel to the orientation of the panel. We also scored the exact duration of each test trial to be able to calculate the proportion of time for each scored behaviour (accounting for some minor deviations in the trials durations). We also intended to score vocalizations but could not do so due to technical problems with the audio recording. A second coder naïve to the hypotheses and theoretical background of the study scored 20% of all trials to assess inter-observer reliability which was acceptable (looking away: K=0.62, N=120; time away: Spearman correlation: r_S_=0.99, N=120, p<0.001; resting: r_S_=0.95, N=120, p<0.001; lateral approach: r_S_=0.97, N=120, p<0.001).

Additionally, we used machine learning to track multiple keypoints of the dogs’ bodies throughout the test trials (Video S2). We used Loopy for the annotation of videos, deep learning based keypoint detection and the 3D reconstruction of keypoints based on video collections of the same trial shot from eight different camera positions (see supplementary material and [36]). For the training of the keypoint detectors, we annotated videos of 38 dogs (36 dogs of the test sample and 2 pilot dogs). We annotated the videos of all eight cameras of one test condition (the test conditions in the annotation sample were counterbalanced). We annotated four keypoints on the dogs’ body: snout, head centre, base of the tail, and the tip of the tail. In total, we annotated 312 videos and for each video, we annotated between one frame per second and one frame every 4 seconds (resulting in 45,686 annotations that were distributed on the four keypoints as follows: snout: 10,446, head centre: 13,141, tail base: 11,245, tail tip: 10,854). Based on these annotations we trained a keypoint detector (with the following setting in Loopy: input network size= 1434 × 1074, stride=8, training iterations=200,000). We used the keypoint detector to predict the 2D keypoint coordinates in all test video collections and reconstructed the 3D coordinates based on these predictions.

Based on the pre-processed data (see supplementary material) we calculated the following variables: whether the dogs visited the interest area around the caregiver within a trial (binary; interest area: 2.44m x 1.68m, the area of the chair plus 1 m or until the wall; see Figure 4A) and the proportion area of the room visited by the dogs. We calculated the proportion area visited based on a function that overlaid a virtual grid consisting of 50 × 50cm cells on the floor plan. The function calculated the proportion of grid cells visited by the dogs. Based on the proportion of tracked data (mean: 99.1 %, min: 78.3%, max: 100%) and in keeping with a previous study [36], we decided to use the head centre keypoint for these analyses.

Additionally, we calculated the 2D angle (based on XY coordinates) between the head-centre— tail-base axis and the tail-base—tail-tip axis (henceforth: tail angle). This angle depended on the tail position relative to the torso but also to the head position relative to the torso though to a smaller extent (because head movements only lead to relatively small deviations of the head-centre—tail-base axis from the torso axis). We therefore used this measure as a proxy for the tail angle with negative values indicating a leftward deflection, 0 indicating no deflection, and positive values a rightward deflection (one dog was excluded due to a docked tail).

#### Analysis

##### Preregistered analyses

Hypotheses, design and the scoring and analysis plan were preregistered (https://aspredicted.org/FHR_1L6). We analysed the duration response variables as proportions (relative to the entire duration of the test trial). The “looking away” response variable was analysed as binary variables (present/absent). For the proportion time response variables, we fitted generalized linear mixed models (GLMM) with beta error distribution (R package glmmTMB [37]). We transformed the data so that they did not comprise the extreme values 0 and 1 [38]. For the binary response variable, we fitted a GLMM with binomial error distribution (R package lme4 [39]). For all GLMMs, we included the following predictor variables: condition and sex as categorical predictors and trial number within condition (1-4), order of conditions (1-3), age (in months) as continuous predictors. Dog ID was included as a random intercept. We also included all possible random slope components. Following our preregistered contingencies plans, we removed the correlation between the random slopes and random intercept for the proportion time response variables due to convergence issues. We z-transformed the covariates and centred all random slope components. Likelihood ratio tests (R function drop1 with argument ‘test’ set to “Chisq”) with p-values smaller than 0.05 were used as criterion to make inferences about fixed effects (see supplementary material for further details).

##### Exploratory analysis of 3D tracking data

We followed the same analysis plan as in preregistered analyses. We analyse the caregiver visits as a binary variable. The proportion area visited variable was analysed using a beta-distributed GLMM (overdispersion parameter: 0.99). We analysed the tail angle by fitting a LMM. We included the same predictor variables as before. Additionally, all models were weighted on the proportion of tracked data.

### Results

The results of the confirmatory and exploratory (3D tracking based) analyses were consistent: the dogs’ behaviour differed significantly between the three conditions. As predicted, they waited more patiently for the food to arrive in the clumsy condition than the teasing condition (shorter periods away from experimenter, less sitting/lying down, less looking away from experimenter, less caregiver visits, and a smaller area of the room explored). In the clumsy condition, the dogs also showed a significant rightward tail deflection compared to the other conditions.

#### Confirmatory analysis

##### Proportion of duration away from experimenter

A GLMM with beta error distribution showed that condition had a significant effect on how long the dogs were away from the experimenter (χ^2^(2)=62.30, p<0.001; Figure 2A). They spent significantly more time away in the blocked than clumsy condition (z=-8.78, p<0.001) or teasing condition (z=-4.78, p<0.001) and they spent more time away in the teasing condition than the clumsy condition (z=-5.07, p<0.001). Besides, the dogs spent significantly more time away with increasing trial number (within condition) and less time away with increasing block number (order of conditions). There were no significant effects of age and sex (Table S1).

**Figure 2.**
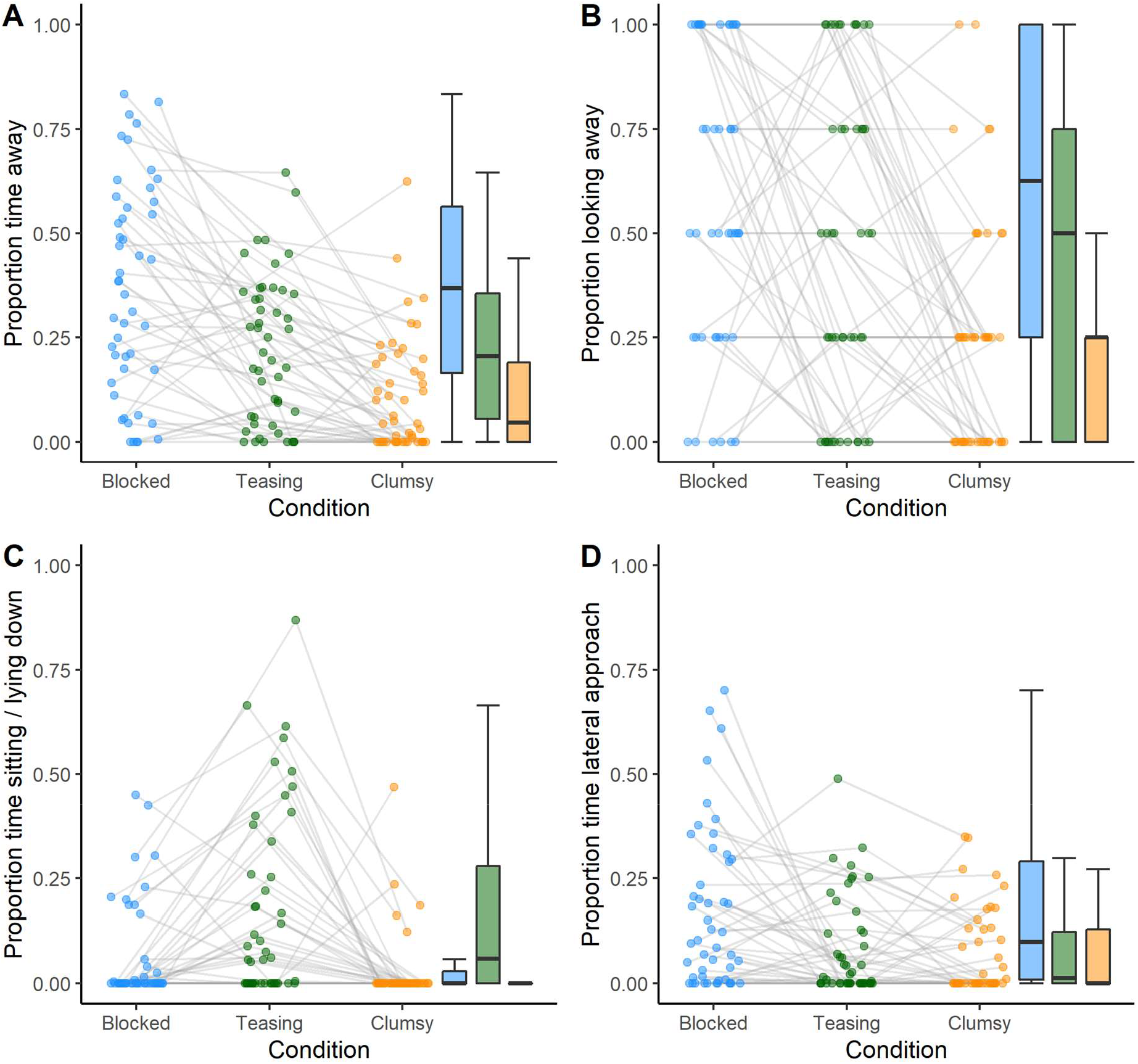
Dot and box plots of dogs’ performance in the test trials. A. Proportion of time away from experimenter, B. Proportion of trials looking away from the experimenter, C. Proportion of time sitting or lying down, D. Proportion of time lateral approach response. The dots represent individual mean values; the grey lines connect the values of the same individuals across conditions. On the right side of each graph, a box plot is shown (blue: blocked condition; green: teasing condition; orange: clumsy condition).

##### Looking away from experimenter

A GLMM with binomial error distribution revealed that condition had a significant effect on the dogs’ likelihood to look away from the experimenter (χ^2^(2)=35.67, p<0.001; Figure 2B). They looked more often away from the experimenter in the blocked than the clumsy (z=-6.20, p<0.001) or teasing condition (z=-2.21, p<0.027) and they looked away more often in the teasing condition compared to the clumsy condition (z=-3.26, p=0.001). Besides, neither trial number, order of conditions, sex or age had a significant effect on the looking away response (Table S2).

##### Proportion duration sitting or lying down

A GLMM with beta error distribution revealed that condition had a significant effect on resting behaviour (χ^2^(2)=16.47, p<0.001; Figure 2C). The dogs spent significantly more time sitting or lying down in the teasing condition than the blocked condition (z=2.96, p=0.003) and the clumsy condition (z=-3.97, p<0.001). The rest response did not differ significantly between the blocked and clumsy condition (z=-1.00, p=0.315). Besides, there was no significant effect of trial number (within condition), block number (order of condition), age or sex on the resting response (Table S3).

##### Lateral approach

A GLMM with beta error distribution revealed that condition had a significant effect on the lateral approach response (χ^2^(2)=17.65, p<0.001; Figure 2D). The dogs spent significantly more time at side of the experimenter in the blocked than in the clumsy (z= -3.98, p<0.001) or teasing condition (z=-3.40, p<0.001) whereas no significant difference was found between the teasing and clumsy condition (z=-0.63, p=0.531). Besides, the dogs spent significantly less time at the side of the experimenter with increasing block number (order of conditions). The trial number, sex or age had no significant effect on the lateral approach response (Table S4).

##### Exploratory analysis based on 3D tracking data

The dogs’ roaming pattern across the four conditions is visualized in Figure 3A (plots of the individual trajectories are available in the associated Github repository: https://github.com/cvoelter/dog_unwilling_unable).

**Figure 3.**
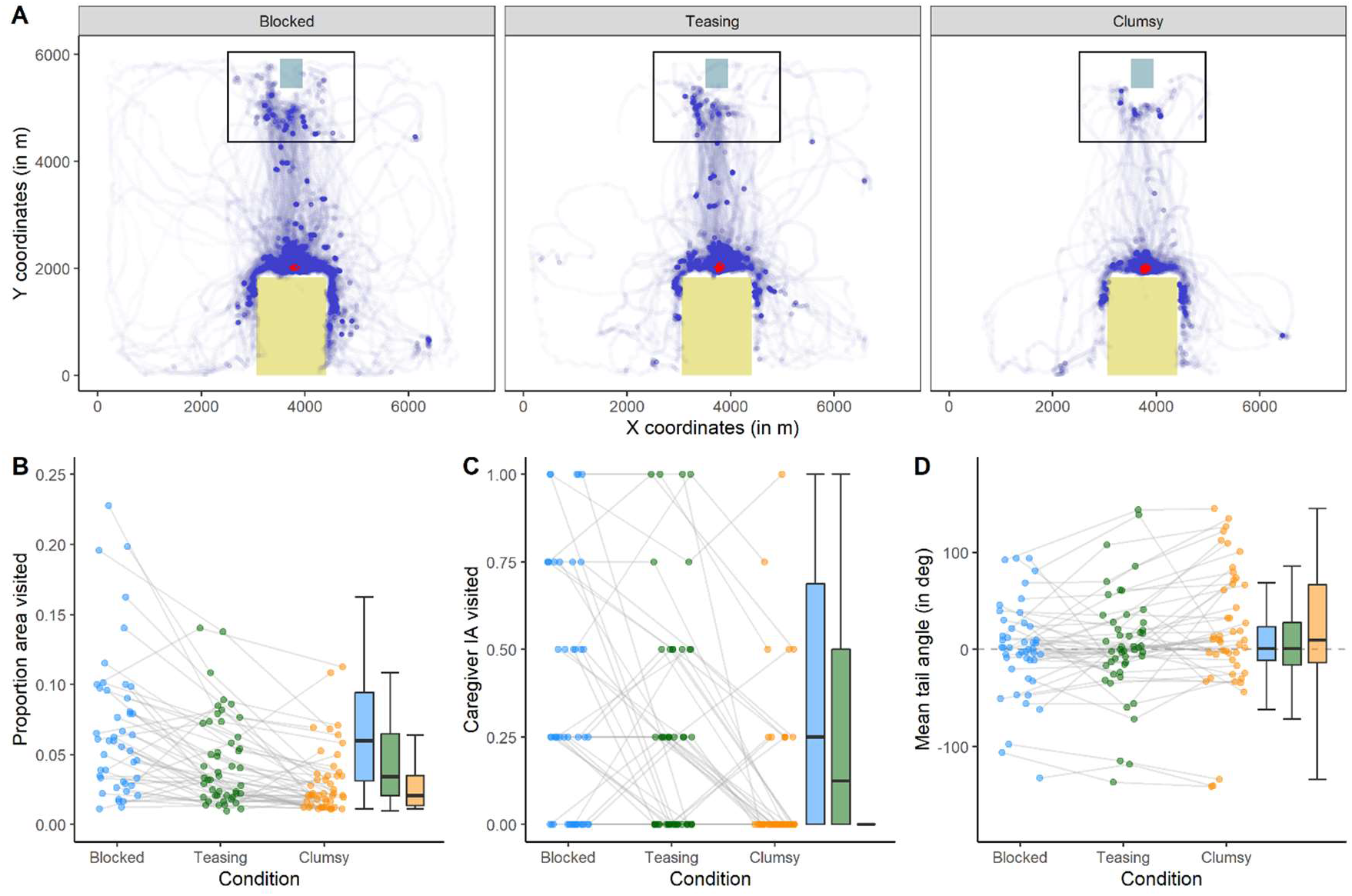
*A*. Plot showing the dogs (N=48) roaming pattern across the four conditions (based on the head-centre keypoint). The purple dots indicate areas visited by the dogs; darker purple areas were visited more frequently. The grey square indicates the location of the caregiver’s chair, the yellow square the location of the fenced area (where the experimenter offered food). The black rectangle indicates the interest area around the caregiver’s location. The red concentric lines show the contours of a 2D kernel density estimation highlighting frequently visited areas. *B*.-*D*. Dot and box plot of dogs’ performance in the test trials based on the 3D tracking data. *B*. Mean proportion of the room area visited. *C*. Mean proportion of trials in which the caregiver interest area was visited. *D*. The angle of the tail relative to the tail-base—head-centre axis (positive values: rightward deflection, negative values: leftward deflection). The dots represent individual mean values; the grey lines connect the values of the same individuals across conditions. On the right of each plot, a box plot shows the median and the quartiles (blue: blocked condition; green: teasing condition; orange: clumsy condition).

##### Proportion of area visited

A GLMM with beta error distribution revealed that condition had a significant effect on the proportion area visited (χ^2^(2) =41.14, *p* < 0.001; Figure 3B): They explored significantly more area in the blocked than clumsy condition (z=-7.94, p<0.001) or teasing condition (z=4.63, p<0.001) and also more in the teasing than the clumsy condition (z=-4.60, p<0.001). Besides, the dogs explored significantly more with increasing trial number (within a block) and they explored significantly less with increasing block number (order of condition). Age and sex had no significant effect on the area explored (Table S5).

##### Caregiver visit (binary)

A binomial GLMM revealed that condition had a significant effect on caregiver visits (χ^2^(2) =25.62, *p* < 0.001; Figure 3C): They visited their caregiver more often in the blocked than the clumsy condition (z=-4.48, p<0.001) but not when compared to the teasing condition (z=-1.90, p=0.057). The also visited the caregiver more often in the teasing condition than the clumsy condition (z=-3.76, p<0.001). Besides, the dogs were more likely to visit their caregiver with increasing trial number. The order of conditions, age and sex had no significant effect on the likelihood of visiting the caregiver (Table S6).

##### Tail angle

A LMM revealed that condition had a significant effect on caregiver visits (χ^2^(2) =8.53, *p=*0.014; Figure 3D): The dogs’ tail was significantly more on the right side of their torso in clumsy condition than in the blocked condition (t=2.92, p=0.006) and the teasing condition (t=2.27, p=0.030). There was no difference between the blocked and teasing condition (t=0.69, p=0.495). Trial number, the order of conditions, age and sex had no significant effect on the tail angle (Table S7).

## Experiment 2

In Experiment 2, our goal was two-fold: to replicate the findings of Experiment 1 and to investigate whether dogs, based on the teasing and clumsy actions, would acquire a preference for a willing but unable experimenter over an unwilling experimenter. As in the previous study, dogs were presented with the unwilling-unable paradigm but this time two different experimenters performed the unwilling-teasing and unable-clumsy conditions.

We predicted (preregistration: osf.io/sjwzm) (1) that dogs would acquire a preference for approaching one of the two experimenters based on the demonstration phase and (2) that they would transfer the intention observed during the demonstration phase of the experiment also to a different context in which the unwilling and unable experimenters acted as informants. Hence, we predicted that they would follow the cues of the unable (clumsy) experimenter more readily compared to those given by the unwilling (teasing) experimenter.

### Methods

#### Subjects

Forty-eight pet dogs of various breeds and mixed-breeds participated in this study (mean age: 89.9 months; range: 17 and 197 months; 32 females and 16 males). Five additional dogs were excluded from the study: two dogs because they entered the fenced area and three further individuals because they did not take the food from the experimenters. The study was discussed and approved by the institutional ethics and animal welfare committee in accordance with GSP guidelines and national legislation (approval number: ETK-132/09/2021).

#### Materials

Same as in Experiment 1. Additionally, we used two green, opaque plastic bowls with paper lids and an occluder made of grey PVC.

#### Design

This experiment included three phases: a warm-up, demonstration and transfer phase (Figure S1). In the warm-up, a neutral experimenter familiarised the dogs with a two-choice pointing task (see supplementary material for details). In the demonstration phase, two different experimenters performed the clumsy and teasing demonstration, respectively. This phase was similar to the clumsy and teasing condition of Experiment 1 but different (female) experimenters (assignment between experimenter ID and conditions were counterbalanced across dogs) performed the conditions. In the transfer phase, we tested whether the dogs had acquired a preference for the willing but unable (clumsy) experimenter over the unwilling experimenter in the demonstration phase. We conducted two transfer tasks: first, a preference task, in which the teasing and clumsy experimenter were just standing passively and the dogs could approach them. Second, an object choice task in which the two experimenters were simultaneously pointing to a different bowl.

In the demonstration phase, the teasing and clumsy trials were presented in blocks in an alternating ABAB block sequence. The order of conditions was counterbalanced across dogs. The dogs received the same number of food rewards from both experimenters during the demonstration phase.

#### Procedure

The demonstration phase was similar to Experiment 1 but two experimenters were inside the fenced area. The two experimenters were sitting back to back on two stools, one experimenter facing and interacting with the dog and the other one facing the wall and looking down. At the beginning of each block and before each test trial, the subject received two motivations trials in which the experimenter successfully passed the food reward (a slice of sausage) to the subject. At the end of each block, there was a short break in which the experimenters swapped positions. In contrast to Experiment 1, the dogs were not called back by the caregiver before each test trial (because there was no blocked condition that would require this) and we made the teasing action more similar to the dropping action of the clumsy condition by moving the hand back and down toward the base of the ramp when moving the food away from the dog. Moreover, the caregiver was asked to wear a blindfold throughout the demonstration phase.

At the beginning of each transfer task, the dogs were sitting in front of the caregiver in an area marked by tape on the floor. The preference test followed the demonstration phase immediately. The two experimenters stepped out of the fenced area and on to marked positions equidistant from the dog. The position of the unwilling/unable experimenter was counterbalanced across dogs. When the experimenters were in position and standing still the caregivers silently counted to three and then released their dog. The caregivers were also instructed to close their eyes when the dogs were released.

The dog could roam freely and approach the experimenters. The preference test took 30 seconds (1 trial). The dog did not receive any food reward during the preference test.

In the subsequent object choice test, the teasing and clumsy experimenter remained on the marked positions (but depending on the counterbalancing scheme they switched sides) while the neutral experimenter placed the material (occluder and two bowls with lids) on the floor. As in the warm-up, the neutral experimenter then showed a food reward to the dog but this time baited both bowls behind the occluder. After the neutral experimenter removed the occluder and slid the bowls to the target positions, she moved behind told the clumsy and teasing experimenter to kneel and point to ensure synchronized movements of both experimenters. The teasing and clumsy experimenter pointed with their ipsilateral arm (right arm on the right side and the left arm on the left side). Both touched the lid of the respective bowl with the index finger. After three seconds, the neutral experimenter told the caregiver to release the dog. The dog could now make one choice and they received the reward from the bowl they approached first (both bowls were baited). After the choice, the caregiver called the dog back. If a dog did not make choice, the caregiver was allowed to encourage the dog verbally after 10 seconds. We conducted eight trials with the position of the clumsy/teasing experimenter counterbalanced across trials. Each trial took a maximum of 30 seconds or until the dogs made a choice.

#### Scoring

For the demonstration phase, we scored the same variables as in Experiment 1 (except for the looking direction). Concerning the preference test, we scored which experimenter (unable/unwilling) the dogs approached first. We scored the first approach once the dogs entered with their snout in one out of two predetermined areas (40×40cm; marked by tape on the floor) around each of the two experimenters. Moreover, we scored how long the dogs spent close to each of the experimenters (in a predetermined area; 90×90cm; marked by tape on the floor). For the object choice test, we scored which cup (unable/unwilling side) the dogs touched first in each trial.

#### Analysis

With respect to the demonstration phase analysis, we followed the same analysis plan as in preregistered analyses of Experiment 1 with the following changes: condition had only two levels (clumsy, teasing), the order of conditions was coded as a factor (teasing-first, clumsy-first), and trial number represented the overall trial number (1-16).

We followed our preregistered analysis plans for the transfer phase (preregistration: osf.io/sjwzm). For the preference test, we analysed the first approach response (coded as binary response: unable 1, unwilling 0). We conducted a binomial test (chance level: 0.5) to evaluate whether the dogs have acquired a significant preference for approaching the unable or unwilling experimenter. We only included dogs in this analysis that approached one of the two experimenters during the 30 seconds of the preference test (N=47). We used a paired-samples t-test to compare the times the dogs spend close to the experimenters (after checking that the difference in the durations was approximately normally distributed). We predicted that based on the demonstrations dogs might have acquired a preference for approaching either the unable experimenter (the dogs might have preferred the person that was more willing to give food) or the unwilling experimenter (the dogs might have attempted to appease or challenge the unwilling experimenter).

For the object choice test, we fitted a GLMM with binomial error structure on the dogs’ first choice within each trial (binary response coded as unable 1, unwilling 0). To evaluate whether dogs have a significant preference for the cup that was cued by the unable demonstrator, we fitted an intercept-only model in which we include the random intercept of subject ID and the random slope of the trial number (1-8; z-transformed to a mean of 0 and a sd of 1). We used the Wald test of the intercept to infer whether their performance deviated significantly (p<0.05) from the hypothetical chance level of 0.5.

## Results

The results of the demonstration phase confirmed the findings of Experiment 1: the dogs reacted differently to the teasing experimenter than the clumsy experimenter (i.e., longer periods away from experimenter and more sitting/lying down in teasing than clumsy condition). In the transfer phase, there was no evidence that the dogs had acquired a significant preference for either of the experimenter neither in the preference test nor in the object choice test.

### Demonstration phase

#### Proportion of duration away from experimenter

A GLMM with beta error distribution revealed that the dogs spent significantly more time away in the teasing than clumsy condition (χ^2^(1)=19.03, p<0.001). There were no significant effects of trial number, order of condition, age and sex (Table S8; Figure S2A).

#### Proportion duration sitting or lying down

A GLMM with beta error distribution provided evidence that dogs spent significantly more time sitting or lying down in the teasing condition than the clumsy condition (χ^2^(1)=18.64, p<0.001). The dogs were also more likely to sit or lie down with increasing trial number (χ^2^(1)=4.11, p=0.043).

Besides, there was no significant effect of the order of condition, age or sex on the resting response (Table S9; Figure S2B).

#### Lateral approach

A GLMM with beta error distribution provided no evidence for a significant effect of condition or the other predictor variables on the lateral approach response (Figure S2C; Table S10).

### Transfer phase

#### Preference test

The dogs had no significant preference for either experimenter in their first approach (mean choice of clumsy experimenter: 0.47, binomial test: p=0.560) or with respect to the time they spent in proximity of the experimenters (t(47)=0.05, p=0.959).

#### Object choice test

The dogs did not show a significant preference for choosing the bowl cued by one of the experimenters (mean choice of bowl cued by clumsy experimenter: 0.46; z=-0.16, p=0.131).

## Discussion

Our study provides robust evidence that dogs’ distinguish between superficially similar human actions that yielded the same outcome but that were clearly different in terms of the underlying intentions: the willingness to transfer food. The dogs behaved as if they understood the underlying intentions, for example, by waiting more patiently for the food to arrive in the clumsy than the teasing condition. We find a number of behavioural markers that evidenced this distinction between the clumsy and teasing conditions: these markers included not only the dogs’ likelihood to stay close to the experimenter but also how often they looked away from the experimenter and how likely they were to sit or lie down. The 3D tracking data show that this distinction can also be found when the behaviour is quantified through machine learning: the proportion of room area visited and the caregiver interest area variables provided complementary measures that reflected whether and to what extent the dogs would stay close to the experimenter throughout the trials.

A new aspect provided by the 3D tracking data was the tail angle measure that yielded evidence for a greater rightward tail deflection in the clumsy condition. Rightward tail asymmetry has been associated with stimuli that should lead to an approach response in dogs in previous studies [40].

Therefore, the rightward tail bias found in the clumsy condition of the current study is supportive of the interpretation that dogs recognized the willingness of the clumsy experimenter to transfer the food.

The dogs’ reaction to the blocked condition was different from the other unable condition, the clumsy condition, in some respect. The dogs approached the experimenter from the side where a wire mesh separated the dogs from the experimenter predominantly in the blocked condition. This seemed to be a response to the change in the physical setup: the closure of the holes in the front panel led them to approach the experimenter from the sides. The lateral barrier was made of a wire mesh that in principle would have allowed transferring the food. The dogs also looked away more often, visited a larger area of the room, and walked back to the caregiver more frequently in the blocked condition than the clumsy condition. Most previous studies using the unwilling-unable paradigm (and that included a clumsy condition) yielded similar results: chimpanzees [4] and capuchin monkey [10] left the experimenter sooner in the blocked and teasing condition than in the clumsy condition (though the clumsy and blocked conditions were not compared statistically in these studies). Trösch et al. [11] found that horses left earlier and spent less time in proximity in the blocked condition than in the clumsy condition. In contrast, Schünemann et al. [35] changed the setup such that the dogs actually could walk around the barrier to approach the experimenter directly. They found that the latency of walking around the barrier was the shortest in the blocked condition, followed by the clumsy condition and then the teasing condition. Given the difference in their setup to our study (and the other instantiations of this paradigm), those results are not directly comparable with ours. Nevertheless, in agreement with our findings, the quicker approach response found by Schünemann et al. [35] suggest that the dogs tried to find another way to get to the human offering the food when the opening was closed.

Another response variable showed a greater similarity in dogs’ response to the two unable conditions (blocked and clumsy) as compared to the unwilling (teasing) condition: the dogs were more likely to sit or lie down in the teasing than the two unable conditions. Schünemann et al. [35] reported a similar pattern but could not confirm this statistically due to a low prevalence of the target behaviour. Schünemann et al. [35] suggested that sitting or lying down might be considered as calming signal that serves to appease the human interaction partner. However, there is only limited support for this interpretation based on the existing literature on calming signals in dogs [41,42].

Another (not mutually exclusive) possibility is that dogs offered one of the most extensively trained behaviours as a begging behaviour particularly when presented with an unwilling experimenter. In line with both of these interpretations, previous research found that dogs sat or lay down also when presented with unknown verbal commands [43,44] or with food they were forbidden to eat by a human watching them [45].

Our results show that dogs (like chimpanzees, capuchin monkeys, Tonkean macaques, grey parrots, and horses) acted *as if* they understood certain human intentions. This, however, does not show that they represent human intentions but just that they distinguish similar actions associated with different intentions. Different cognitive abilities might support this distinction ranging from associative learning to (some kind of) theory of mind, with even a limited concept of intention being considered as a central part of theory of mind [46]. These possibilities are of course not mutually exclusive and they apply likewise to the other species that were tested in this task (including human infants). In the case of pet dogs, given their daily exposure to human intentional actions, it is possible that they had learnt to distinguish between teasing actions and clumsy dropping actions prior to the study. Dropping food might often result in the dogs obtaining the food in their daily lives. This expectation to obtain the food possibly could explain their greater patience following the clumsy demonstration. The fact that the dogs could not directly approach the dropped food (as in [35]) alleviates this caveat to some extent.

In the context of a growing literature on dogs’ comprehension of referential communicative signals [17,21,34,44,47,48], perspective taking abilities [25,27], goal [28,33] and (false) belief attribution [26] it appears also possible that the current findings reflect a more general and flexible cognitive mechanism allowing dogs to predict human behaviour based on inferred current intentions. A prediction that one could derive from a genuine intention-reading ability is that dogs might expect a human with a certain intention to behave consistently across different contexts. If so, dogs can be expected to avoid an unwilling experimenter in a different situation in which the dogs choose to interact with one of the experimenters.

Therefore, we investigated in Experiment 2 whether the unwilling-unable demonstrations would affect any subsequent interactions between experimenter and subject. However, we found no such evidence: in Experiment 2, the dogs’ experience with an unwilling (teasing) and unable (clumsy) experimenter did not lead them to acquire a significant preference for either of the experimenters. As always, there are multiple possible explanations for such negative findings: for example, it might be that the demonstration phase was too short for the dogs to acquire a preference, any preference the dogs might have was washed out by the intermixed motivation trials, or that the dogs did not generalize their experience to a different situation. These possibilities also highlight opportunities for future research, for instance, by increasing the amount of experience with the clumsy and teasing experimenters, omitting motivation trials (or performing the motivation trials via a third, neutral experimenter), and testing the dogs’ preferences for any of the experimenters without any change in context. Finally, differing motivations between individuals might have led to a null result at the group level. It is conceivable that some dogs preferred approaching the teasing experimenter as appeasement response whereas other dogs preferred approaching the clumsy experimenter as a strategy to gain more food. Administering more trials to allow for individual level analyses and studying how personality traits affects the dogs’ choices might shed more light on this possibility.

In summary, the current study provides robust evidence across two pre-registered studies that pet dogs distinguished between similar human actions leading to the same outcome and act as if they were sensitive to different human intentions. Machine learning based 3D tracking confirmed this distinction and provided further evidence that the dogs interpreted the clumsy (willing but unable) experimenter in a more positive manner (evidenced by the rightward bias in their tail deflection). Our study highlights pet dogs’ sensitivity to subtle differences in human actions and shows how they adjust their behaviour accordingly. How exactly they acquire such behaviour- or intention-reading abilities will be an exciting topic for future research.

## Acknowledgements

We thank Karin Bayer for administrative support, Max Hofbauer and John Stowers (Loopbio), Wolfgang Berger, and Peter Füreder for technical support, and Roger Mundry for his help with the data processing and analysis. Furthermore, we are grateful to the dog caregivers for participating with their dogs in this study.

This project was supported by the Austrian Science Fund (FWF) through project W1262-B29 to LH.

## Authors’ contributions

C.J.V. conceptualization, supervision, data curation, formal analysis, methodology, project administration, visualization, writing-original draft, writing-review and editing; L.L. conceptualization, supervision, methodology, writing-review and editing; M.G.G.M.S. methodology, data collection, writing-review and editing; C.F.R. methodology, data collection, writing-review and editing; K.G. methodology, data collection, writing-review and editing; M.S. methodology, data collection, writing-review and editing; I.D. annotation, methodology; L.H. conceptualization, funding acquisition, methodology, resources, writing-review and editing. All authors approved the final version of the manuscript.

## Data availability statement

The data and R scripts associated with this manuscript are available on GitHub: https://github.com/cvoelter/dog_unwilling_unable

