## Supplementary text and figures for "Unwilling or Unable? Using 3D tracking to evaluate dogs’ reactions to differing human intentions"

### Experiment 1: supplementary methods

#### Camera setup

The first camera system consisted of four HD cameras (Panasonic HC-V777) mounted to the walls of the testing room that filmed the test session from different perspectives. The second camera system consisted of eight 4K cameras (Theia Technologies ML410M, 1/1.7", 4-10mm, F1.4) mounted below the ceiling in every corner of the room and in the centre of each wall. The former camera system was used for the manual scoring; the latter was used for the 3D body tracking.

#### Pre-processing of 3D tracking data

The 3D data were pre-processed as follows: (1) we removed values deemed unrealistic (coordinates outside of the room and coordinates higher than 1.5 m from the ground), (2) we determined the average coordinates across the four key points for each video frame and removed individual key points whose coordinates deviated more than 1 m from the average coordinate, (3) we excluded coordinates that deviated more than 1.5 m in any dimension from the previous frame (video frame rate: 25 fps). We conducted a linear interpolation to fill in missing data. Then we filtered out the data that deviated more than 20 centimetres in any direction from one frame to the next and conducted another linear interpolation. We repeated the last step (filtering followed by interpolation) three more times. Finally, we calculated a rolling average with a window size of 3 frames to smoothen the trajectories.

#### Analysis: assumption checks

To evaluate the model stability, we removed one subject at a time and compared the resulting estimates. This procedure revealed the models stable with respect to fixed effects. For the beta models, we also checked for overdispersion, which was not an issue (dispersion parameter time away: 0.75; sitting/lying down: 1.23). Additionally, we checked for collinearity among the predictor variables, which was no issue either (maximal Variance Inflation Factor for all models: 1.00).

For the tail angle analysis, wwe checked the assumptions of homogeneous and normally distributed residuals by visually inspecting the residuals against fitted data and the Q-Q plot of the residuals; we found no obvious violations of the assumptions. Collinearity was no issue (max. VIF: 1.00).

### Experiment 2: supplementary methods

#### Procedure: warm-up

In the warm-up, the dog sat in front of the caregiver in a rectangular area marked on the floor. The neutral experimenter hid food in one of two bowls outside of the dog's view behind an occluder, closed the bowl by means of a paper lid, removed the occluder and slid the bowl simultaneously to two markings on the floor, equidistant from the dog. The experimenter then pointed to (with the ipsilateral arm) and looked at the baited cup. After three seconds, while holding that pose, the experimenter said “OK” and the caregiver released the dog who could choose one of the cups. If the dog chose the cup correctly, the dog was allowed to eat the treat and the procedure was repeated on the other side. If the dog chose incorrectly (empty bowl), the neutral experimenter took away the baited bowl, the caregiver called the dog back to the starting position and the procedure was repeated once on the same side, before it was performed on the other side. The goal of the warm-up was to familiarize the dogs with the task. In particular, the dogs should experience that only one cup contained food and that they were allowed to make only one choice. After the warm-up, the neutral experimenter removed the bowls and the occluder and left the room.

**Setup**

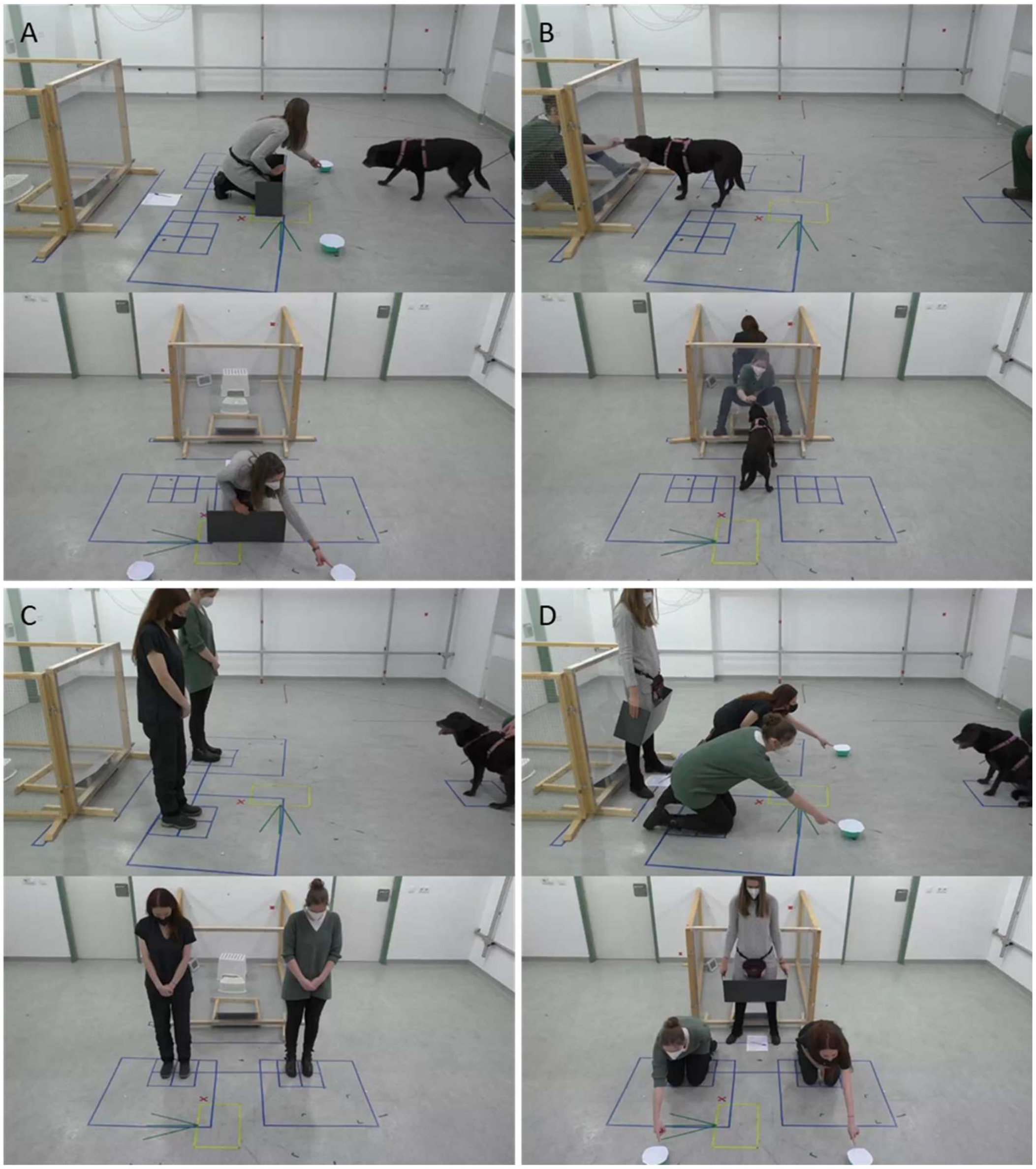


*Figure S1 Screenshots of the different experimental phases. A) Warm-up phase: the neutral experimenter familiarizes the dogs with the object choice test; B) Demonstration phase with the clumsy and teasing experimenters in the fenced area; C) Preference test; D) Pointing test*. *Informed consent was obtained for publication of identifying images.*

**Analysis: assumption checks**

The models were stable with respect to fixed effects. For the beta models, we also checked for overdispersion which only provided some indication for overdispersion in the case of the lateral approach response (dispersion parameter time away: 0.72; sitting/lying down: 0.69, lateral approach: 1.39). Additionally, we checked for collinearity among the predictor variables, which was no issue either (maximal Variance Inflation Factor for all models: 1.09).

Results

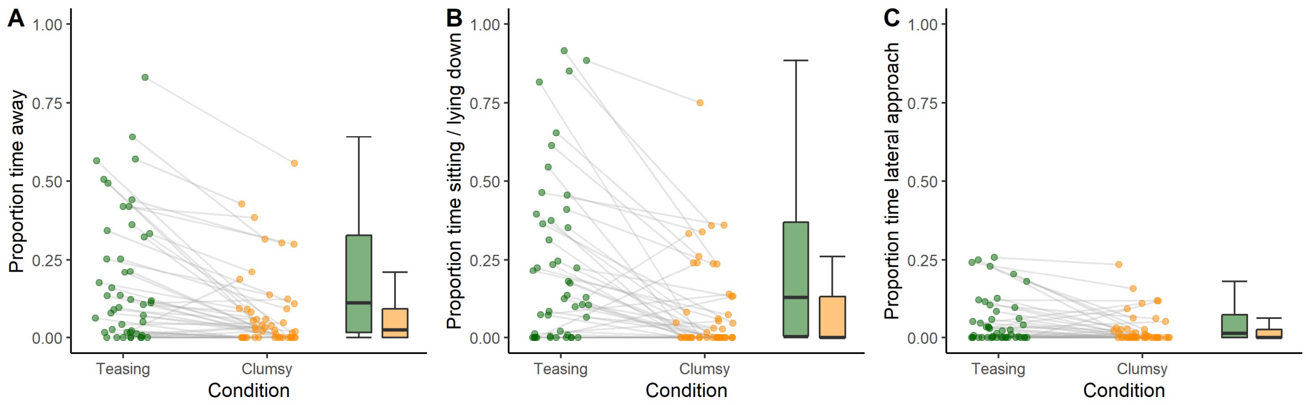


*Figure S2* Dot and box plots of dogs' performance in the demonstration phase of Experiment 2. A. Proportion of time away from experimenter, B. Proportion of time sitting or lying down, C. Proportion of time of lateral approach response. The dots represent individual mean values; the grey lines connect the values of the same individuals across conditions. On the right side of each graph, a box plot is shown (green: teasing condition; orange: clumsy condition).

*Table S1 Results of the GLMM 01 with duration away as response variable*

| | Estimate | SE | LowerCI | UpperCI | $\chi^2$ | df | P |
|---|---|---|---|---|---|---|---|
| (Intercept) | -0.93 | 0.15 | -1.23 | -0.65 | | | |
| Condition - clumsy[1] | -1.16 | 0.13 | -1.43 | -0.91 | 62.30 | 2 | <0.001 |
| Condition - teasing[1] | -0.59 | 0.12 | -0.84 | -0.36 | | | |
| Trial | 0.24 | 0.05 | 0.14 | 0.33 | 25.32 | 1 | <0.001 |
| Order | -0.14 | 0.06 | -0.26 | -0.03 | 5.77 | 1 | 0.016 |
| Age | 0.05 | 0.1 | -0.16 | 0.25 | 0.21 | 1 | 0.646 |
| Sex[2] | 0.24 | 0.21 | -0.19 | 0.62 | 1.24 | 1 | 0.266 |

Notes: Reference category: [1]blocked, [2]female; all covariates were z-transformed to a mean of 0 and an sd of 1.

*Table S2 Results of the GLMM 02 with looking away as response variable*

| | Estimate | SE | LowerCI | UpperCI | $\chi^2$ | df | P |
|---|---|---|---|---|---|---|---|
| (Intercept) | 0.88 | 0.5 | 0.02 | 2.07 | | | |
| Condition - clumsy[1] | -2.77 | 0.45 | -4.10 | -2.12 | 35.67 | 2 | <0.001 |
| Condition - teasing[1] | -1.22 | 0.55 | -2.47 | -0.23 | | | |
| Trial | 0.83 | 0.15 | 0.56 | 1.2 | 28.75 | 1 | <0.001 |
| Order | -0.36 | 0.21 | -0.83 | 0.01 | 2.98 | 1 | 0.084 |
| Age | 0.42 | 0.28 | -0.20 | 1.05 | 2.28 | 1 | 0.131 |
| Sex[2] | 0.11 | 0.62 | -1.09 | 1.34 | 0.03 | 1 | 0.856 |

Notes: Reference category: [1]blocked, [2]female; all covariates we z-transformed.

*Table S3 Results of the GLMM 03 with sitting / lying down as response variable*

| | Estimate | SE | LowerCI | UpperCI | $\chi^2$ | df | P |
|---|---|---|---|---|---|---|---|
| (Intercept) | -2.22 | 0.11 | -2.44 | -2.02 | | | |
| Condition - clumsy[1] | -0.11 | 0.11 | -0.30 | 0.10 | 16.47 | 2 | <0.001 |
| Condition - teasing[1] | 0.33 | 0.11 | 0.10 | 0.54 | | | |
| Trial | 0.01 | 0.05 | -0.08 | 0.10 | 0.05 | 1 | 0.829 |
| Order | -0.01 | 0.04 | -0.09 | 0.08 | 0.05 | 1 | 0.821 |
| Age | -0.01 | 0.04 | -0.10 | 0.08 | 0.02 | 1 | 0.886 |
| Sex[2] | <0.01 | 0.09 | -0.18 | 0.19 | <0.01 | 1 | 0.999 |

Notes: Reference category: [1]blocked, [2]female; all covariates we z-transformed.

*Table S4 Results of the GLMM 05 with lateral approach as response variable*

| | Estimate | SE | LowerCI | UpperCI | $\chi^2$ | df | P |
|---|---|---|---|---|---|---|---|
| (Intercept) | -1.91 | 0.11 | -2.13 | -1.69 | | | |
| Condition - clumsy[1] | -0.44 | 0.11 | -0.65 | -0.22 | 17.65 | 2 | <0.001 |
| Condition - teasing[1] | -0.38 | 0.11 | -0.59 | -0.16 | | | |
| Trial | 0.02 | 0.04 | -0.07 | 0.11 | 0.21 | 1 | 0.645 |
| Order | -0.18 | 0.05 | -0.27 | -0.08 | 14.88 | 1 | <0.001 |
| Age | -0.01 | 0.06 | -0.14 | 0.11 | 0.03 | 1 | 0.865 |
| Sex[2] | 0.17 | 0.13 | -0.09 | 0.42 | 1.68 | 1 | 0.195 |

Notes: Reference category: [1]blocked, [2]female; All covariates we z-transformed.

*Table S5 Results of the GLMM 06 with proportion of explored area as response variable*

| | Estimate | SE | $\chi^2$ | df | P |
|---|---|---|---|---|---|
| (Intercept) | -2.85 | 0.10 | | | |
| Condition - clumsy[1] | -0.71 | 0.09 | 41.14 | 2 | <0.001 |
| Condition - teasing[1] | -0.38 | 0.08 | | | |
| Trial | 0.1 | 0.02 | 17.31 | 1 | <0.001 |
| Order | -0.11 | 0.03 | 11.17 | 1 | 0.001 |
| Age | 0.02 | 0.07 | 0.06 | 1 | 0.799 |
| Sex[2] | 0.18 | 0.15 | 1.33 | 1 | 0.248 |

Notes: Reference category: [1]blocked, [2]female; all covariates were z-transformed to a mean of 0 and an sd of 1.

*Table S6 Results of the GLMM 07 with caregiver IA visited as response variable*

| | Estimate | SE | LowerCI | UpperCI | $\chi^2$ | df | P |
|---|---|---|---|---|---|---|---|
| (Intercept) | -1.25 | 0.49 | -2.35 | -0.36 | | | |
| Condition - clumsy[1] | -2.75 | 0.61 | -4.46 | -1.78 | 25.62 | 2 | <0.001 |
| Condition - teasing[1] | -0.72 | 0.38 | -1.55 | 0 | | | |
| Trial | 0.72 | 0.15 | 0.46 | 1.09 | 24.4 | 1 | <0.001 |
| Order | 0.11 | 0.18 | -0.23 | 0.47 | 0.39 | 1 | 0.533 |
| Age | 0.28 | 0.33 | -0.41 | 0.98 | 0.73 | 1 | 0.393 |
| Sex[2] | 0.16 | 0.7 | -1.25 | 1.53 | 0.05 | 1 | 0.824 |

Notes: Reference category: [1]blocked, [2]female; all covariates were z-transformed to a mean of 0 and an sd of 1.

*Table S7 Results of the GLMM 08 with tail angle visited as response variable*

| | Estimate | SE | LowerCI | UpperCI | $\chi^2$ | df | P |
|---|---|---|---|---|---|---|---|
| (Intercept) | 179.49 | 9.66 | 160.22 | 198.53 | | | |
| Condition - clumsy[1] | 17.62 | 5.92 | 4.01 | 30.54 | 8.53 | 2 | 0.014 |
| Condition - teasing[1] | 4.02 | 5.7 | -8.82 | 17.66 | | | |
| Trial | -1.44 | 2.39 | -6.84 | 4.02 | 0.36 | 1 | 0.548 |
| Order | 3.02 | 2.38 | -2.1 | 8.5 | 1.52 | 1 | 0.218 |
| Age | -7.48 | 7.24 | -21.45 | 6.25 | 1.06 | 1 | 0.304 |
| Sex[2] | -0.15 | 15.06 | -29.97 | 27.33 | <0.001 | 1 | 0.992 |

Notes: Reference category: [1]blocked, [2]female; all covariates were z-transformed to a mean of 0 and an sd of 1.

*Table S8 Results of the GLMM 09 with duration away as response variable*

| | Estimate | SE | LowerCI | UpperCI | $\chi^2$ | df | P |
|---|---|---|---|---|---|---|---|
| (Intercept) | -2.26 | 0.18 | -2.61 | -1.93 | | | |
| Condition - teasing[1] | 0.45 | 0.09 | 0.25 | 0.64 | 19.03 | 1 | <0.001 |
| Trial | 0.07 | 0.05 | -0.02 | 0.16 | 2.03 | 1 | 0.155 |
| Order[2] | 0.14 | 0.22 | -0.29 | 0.54 | 0.38 | 1 | 0.538 |
| Age | 0.07 | 0.11 | -0.14 | 0.30 | 0.36 | 1 | 0.546 |
| Sex[3] | -0.06 | 0.24 | -0.48 | 0.41 | 0.07 | 1 | 0.796 |

Notes: Reference category: [1]clumsy, [2]clumsy-first, [3]female; all covariates were z-transformed to a mean of 0 and an sd of 1.

*Table S9 Results of the GLMM 10 with sitting / lying down as response variable*

| | Estimate | SE | LowerCI | UpperCI | $\chi 2$ | df | P |
|---|---|---|---|---|---|---|---|
| (Intercept) | -1.86 | 0.17 | -2.19 | -1.53 | | | |
| Condition - teasing[1] | 0.56 | 0.12 | 0.33 | 0.80 | 18.64 | 1 | <0.001 |
| Trial | 0.09 | 0.04 | 0.00 | 0.18 | 4.11 | 1 | 0.043 |
| Order[2] | 0.08 | 0.19 | -0.33 | 0.50 | 0.16 | 1 | 0.692 |
| Age | 0.19 | 0.1 | 0.00 | 0.39 | 3.41 | 1 | 0.065 |
| Sex[3] | -0.13 | 0.21 | -0.54 | 0.27 | 0.38 | 1 | 0.537 |

Notes: Reference category: [1]clumsy, [2]clumsy-first, [3]female; all covariates we z-transformed.

*Table S10 Results of the GLMM 11 with lateral approach as response variable*

| | Estimate | SE | LowerCI | UpperCI | $\chi2$ | df | P |
|---|---|---|---|---|---|---|---|
| (Intercept) | -2.94 | 0.09 | -3.15 | -2.77 | | | |
| Condition - teasing[1] | 0.12 | 0.07 | -0.02 | 0.28 | 2.8 | 1 | 0.094 |
| Trial | 0.01 | 0.04 | -0.06 | 0.08 | 0.03 | 1 | 0.855 |
| Order[2] | 0.02 | 0.07 | -0.13 | 0.18 | 0.08 | 1 | 0.781 |
| Age | -0.06 | 0.04 | -0.13 | 0.01 | 2.61 | 1 | 0.106 |
| Sex[3] | -0.03 | 0.08 | -0.18 | 0.13 | 0.12 | 1 | 0.733 |

Notes: Reference category: [1]clumsy, [2]clumsy-first, [3]female; all covariates we z-transformed.
